# Temporal dynamics of visual representations in the infant brain

**DOI:** 10.1101/2020.02.26.947911

**Authors:** Laurie Bayet, Benjamin D. Zinszer, Emily Reilly, Julia K. Cataldo, Zoe Pruitt, Radoslaw M. Cichy, Charles A. Nelson, Richard N. Aslin

## Abstract

Tools from computational neuroscience have facilitated the investigation of the neural correlates of mental representations. However, access to the representational content of neural activations early in life has remained limited. We asked whether patterns of neural activity elicited by complex visual stimuli (animals, human body) could be decoded from EEG data gathered from 12-15-month-old infants and adult controls. We assessed pairwise classification accuracy at each time-point after stimulus onset, for individual infants and adults. Classification accuracies rose above chance in both groups, within 500 ms. In contrast to adults, neural representations in infants were not linearly separable across visual domains. Representations were similar within, but not across, age groups. These findings suggest a developmental reorganization of visual representations between the second year of life and adulthood and provide a promising proof-of-concept for the feasibility of decoding EEG data within-subject to assess how the infant brain dynamically represents visual objects.

## 1. Introduction

A key question in developmental cognitive science concerns the content and properties of neural representations in preverbal infants: How do these representations change with brain maturation and experience? Because infants cannot explicitly report on their own representations, developmental scientists implicitly probe representations by measuring looking times and other behaviors (Aslin, 2007). These behavioral paradigms are limited to testing a few stimulus contrasts, given the short duration of cooperativity in infants (Aslin & Fiser, 2005). Neural measures such as functional near-infrared spectroscopy (fNIRS) and electroencephalography (EEG) additionally reveal spatial (Lloyd-Fox, Blasi, & Elwell, 2010) and temporal (Csibra, Kushnerenko, & Grossmann, 2008) differences in brain activation in response to experimental conditions, demonstrating already intricate functional specialization in the infant human brain (Dehaene-Lambertz & Spelke, 2015). Like most behavioral measures, these neural activations typically consist of group averages in response to two or three stimulus conditions. Another limitation of these studies is that they focus on differences in the amplitude or timing of a neural signature (e.g., an average evoked component) rather than how reliably each signature maps onto a specific stimulus. Thus, the field of developmental cognitive neuroscience has remained largely based on studies of average activations, leaving open the question of the information that is represented in these neural signals, i.e. neural representations. Multivariate pattern analysis (MVPA) addresses this gap by asking whether one may extract information about relevant aspects of the stimuli presented from patterns of neural activations with higher than chance accuracy (that is, classification), which is then taken to suggest that neural representations support the discrimination of these stimuli (Haxby, Connolly, & Guntupalli, 2014; Haxby et al., 2001; Hung, Kreiman, Poggio, & DiCarlo, 2005; Isik, Meyers, Leibo, & Poggio, 2014; King et al., 2018; Meyers, Freedman, Kreiman, Miller, & Poggio, 2008; Norman, Polyn, Detre, & Haxby, 2006). Here we ask whether patterns of neural activation support the discrimination of stimuli that have been presented to a given infant across trials.

Very few studies have used information-focused methods, such as MVPA, to address this question in infants. Two recent studies have used group-level MVPA-related methods to uncover representations of a small number of stimulus classes in awake infants using near infra-red spectroscopy (fNIRS; Emberson, Zinszer, Raizada, & Aslin, 2017) and fMRI: Deen et al., 2017). These studies provided an early demonstration that information-based analysis methods can map the geometry of neural representations in infant visual cortices at the group level. Here, we extend that work by using time-resolved, within-subject MVPA (Groen, Ghebreab, Lamme, & Scholte, 2012; Grootswagers, Wardle, & Carlson, 2017; Kaneshiro, Perreau Guimaraes, Kim, Norcia, & Suppes, 2015; Kietzmann, Gert, Tong, & König, 2017; Kong, Kaneshiro, Yamins, & Norcia, 2020) of EEG data to reveal the temporal dynamics of neural representations at the level of individual infants.

We focus on two classes of visual stimuli that are highly familiar to infants and easily discriminable based on behavioral studies – animals and parts of the body. Adults identify visual objects quickly and efficiently: the organization of neural representations of visual objects according to specific domains (e.g., animal versus human body) is evident as early as 100-200 ms post-onset in adults, reflecting robust specificity of neural responses along the ventral stream (Cichy, Pantazis, & Oliva, 2014; Isik et al., 2014). Infants already exhibit some functional specificity in their neural responses to visual objects, such as strikingly domain-specific cortical activations to faces by 2-3 months (de Haan, Pascalis, & Johnson, 2002; de Heering & Rossion, 2015; Deen et al., 2017; Farzin, Hou, & Norcia, 2012; Halit, Csibra, Volein, & Johnson, 2004; Tzourio-Mazoyer et al., 2002), and perhaps to animate versus inanimate objects by 7-8-months (Jeschonek, Marinovic, Hoehl, Elsner, & Pauen, 2010; Peykarjou, Wissner, & Pauen, 2017). By the end of the first year of life, infants also demonstrate sensitivity to some visual categories within the animate domain, including perceptual narrowing to human vs. non-human faces (de Haan et al., 2002; Peykarjou, Pauen, & Hoehl, 2014), and categorization of human vs. animal body images in behavioral and oddball ERP tasks (Marinović, Hoehl, & Pauen, 2014; Oakes, Plumert, Lansink, & Merryman, 1996; Pauen, 2000; Quinn & Eimas, 1998). However, whether the specificity of visual activations is sufficient to support robust, automatic, and fast representations of visual objects and their categorical domains (e.g. human body parts or animals) in infancy remains unknown. We sought to establish the feasibility of time-resolved MVPA of EEG data in 12-to 15-month-old infants, and in so doing, to probe for the first time the dynamics of neural representations of two types of animate visual objects using machine-learning based classification techniques.

## 2. Materials and Methods

### 2.1. Participants

Within-subject multivariate pattern classification analyses require that each included participant contribute sufficient data to train classifiers, which is challenging to achieve with infants (Emberson et al., 2017). Thus, we restricted analyses to individual participants who contributed at least 50% valid trials compared to the maximum possible number of trials (infants: at least 80 valid trials total out of a maximum of 160, or an average of 10 valid trials per condition; adults: at least 128 valid trials out of a maximum of 256, or an average of 16 valid trials per condition; range: infants 9-19, adults 15-32). The final sample was comprised of 10 12-to 15-month-olds (6 girls, mean age 435.00 ± 20.40 days), and 8 young adults. An additional 12 12-to 15-month-old infants (5 girls, mean age 426.92 ± 21.16), and 4 young adults completed the study but were excluded due to contributing too few valid trials (11 infants, 1 adult), having more than 20% of channels identified as noisy by PrepPipeline (Bigdely-Shamlo, Mullen, Kothe, Su, & Robbins, 2015) during preprocessing of the raw continuous EEG (3 adults), or refusal to wear the EEG net (1 infant). Ages of included versus excluded infants did not significantly differ (two-sample *t*-test, *p* > 0.37). A subset of participating families also completed the English MacArthur Communicative Development Inventories: Words and Gestures (CDI, Infant form) questionnaire (Fenson, 2002); raw CDI scores from this subsample are reported in **Supplementary Table 1**, along with group average scores on word items corresponding to the visual stimuli used in the current study (e.g., “cat”). Briefly, all infants were reported by their caregiver to understand at least one of the CDI words that could describe the visual stimuli used in this study (e.g., most commonly, “dog”, “nose”, “mouth”, “foot”, and/or “cat”); a majority of infants were reported to produce at least one of these words (most commonly, “dog”; see **Supplementary Table 1**). Thus, infants were generally familiar with the depicted objects, as reported by their caregivers. Adult participants, and infant participants’ caregivers, provided written informed consent before the study, which was approved by the Institutional Review Board of the University of Rochester and Boston Children’s Hospital, respectively.

### 2.2. Stimuli

Stimuli were color images of 4 animals (cat, dog, bunny, teddy bear) and 4 parts of the body (hand, foot, mouth, or nose). Pictures were cropped, placed on a uniform gray background, and displayed with a visual angle of roughly 19° by 19° for infants or 8° by 8° for adults. See **Supplementary Figure 1** and Bergelson & Swingley (2012) for details on these stimuli.

### 2.3. Paradigm

Stimuli were presented in random order for 500 ms with a jittered ITI of 1-1.5 s. For infants, stimuli were presented using EPRIME (Schneider, Eschman, & Zuccolotto, 2002) for up to 20 repeated blocks corresponding to a maximum of 160 trials. To avoid presenting a stimulus when the infant was not attending to the screen, presentation of each stimulus was triggered manually by the experimenter watching a live video feedback from an adjacent room. For adults, stimuli were presented using MATLAB and Psychtoolbox (Brainard, 1997) for up to 32 repeated blocks corresponding to a maximum of 256 trials. Adults also saw additional trials corresponding to inanimate stimuli (food and clothing items) which were not seen by infants and thus were not included in the current analyses.

### 2.4. EEG recordings

Infants’ EEG data were recorded at 1000Hz from 128-channels EGI High-Density Geodesic Sensor Nets, referenced online to Cz. A video of children’s behavior while looking at the screen was recorded simultaneously and coded offline. Adults’ EEG data were recorded at 1000Hz from 32-channels BrainVision actiChamp caps, referenced online to the left ear.

### 2.5. EEG preprocessing

Raw continuous EEG signals were processed through the PrepPipeline toolbox (Bigdely-Shamlo et al., 2015) for noisy channel detection and interpolation, robust average-reference, and line-noise removal. Resulting continuous signals were filtered to 0.2-200 Hz using a Butterworth design as implemented in ERPLab’s “pop_basicfilter” function (Lopez-Calderon & Luck, 2014), and further processed using EEGLAB and custom functions (Delorme & Makeig, 2004) as described below. Filtered continuous signals were smoothed using a 20-ms running average, epoched from −50 to 500 ms relative to stimulus onset, and baseline corrected based on the period from −50 to 0 ms relative to stimulus onset. A relatively short epoch duration was chosen in order to maximize the number of valid, artifact-free trials available for multivariate pattern analyses. For infants, outer rim channels were excluded from further analysis to reduce the number of classification features (see **Figure 3** for maps of included channels). Because multivariate pattern analyses rely on patterns of activity across channels, trials were excluded if signals in any scalp channel exceeded ± 150 μV for infants or ± 80 μV for adults. For infants, videos were additionally coded offline to exclude trials when the child stopped looking at the screen during stimulus presentation for any reason (e.g., due to eye movement, head movement, or blink). For adults, individual epochs were additionally excluded if signals in Fp1/2 exceeded ± 60 μV (suspected blink artefact) or signals in horizontal EOGs exceeded ± 40 μV (suspected eye movement artefact). Horizontal EOGs were excluded from further analysis.

### 2.6. Univariate ERP analyses

We report average event-related potentials (ERPs) in each condition and group, using the following regions and time-windows of interest as described in the literature for comparable paradigms: for infants (De Haan, 2013), ERP analyses focused on the N290 (15 occipital electrodes around Oz, O1 and O2; 190-300ms), P400 (18 occipitotemporal electrodes around T5/P7 and T6/P8; 300-500 ms), and NC (16 central electrodes comprising Cz, C3 and C4; 300-500 ms); for adults, ERP analyses focused on the N170 (T5/P7 and T6/P8; 150–190 ms; e.g. Balas & Koldewyn, 2013). ERP analyses were performed in MATLAB.

### 2.7. Multivariate EEG analyses

Time-resolved, within-subject multivariate pattern analysis was performed. An advantage of within-subject classification methods is that classification can leverage individual idiosyncrasies in neural patterns in each individual participant because classifiers do not need to generalize across different participants. Multivariate pairwise classification analyses were conducted using linear SVMs as implemented in libsvm 3.11 (Chang & Lin, 2011) for MATLAB, with 4-fold cross-validation and pseudo-averaging of individual trials within each fold (Grootswagers et al., 2017). That is, to train a given classifier for a given participant, for a given pair of stimuli, and at a given time-point, trials from each of the two stimulus conditions of the pair (e.g., cat vs dog) were randomly re-ordered (permuted) and separated into 4 folds (i.e., quartiles of trials for that stimulus pair). Then, for each of these two conditions, trials from each of the 4 folds were separately averaged to yield 4 pseudo-trials for each of the two conditions, i.e., one pseudo-trial per fold and condition (Grootswagers et al., 2017; Isik et al., 2014). The first 3 of these 4 pseudo-trials were used for training the classifier, while the remaining pseudo-trial was used for testing (i.e., 4-folds cross-validation). As there could be variations in the exact number of valid trials per condition and participant, the exact number of trials that were averaged to form each pseudo-trial could vary. The procedure of re-ordering trials, separating into folds, and training and testing classifiers at every time point was repeated 200 times; classification accuracies are averaged over these instances to yield more stable estimates. Multivariate patterns of channel baseline-normalized amplitude z-scores at each trial were used as features, i.e., for each trial and each channel, voltage amplitudes were z-scored based on the average and standard deviation of voltage amplitudes during the baseline period for that channel and trial. Trials were classified according to stimulus conditions, considering each time-point post-onset and each possible pair of stimulus conditions (e.g., cat vs dog) independently, resulting in a time-series of pairwise stimulus classification accuracy. Thus, the theoretical chance level was 50%. For visualization purposes, classifier weights were transformed back into multivariate activation patterns using the formula proposed in Equation 6 of Haufe et al. (2014). Temporal generalization analyzes (King & Dehaene, 2014) were additionally conducted, by which classifiers are trained on a given time-point post stimulus onset on the training set (e.g. +20 ms post-onset) and trained on another time point on the test set (e.g. +40 ms post onset). Thus, the temporal generalization classifier must generalize not only to unseen data from the same participant, but to a different processing stage post-stimulus; the method thus allows for examining whether neural representations of stimuli are sustained or reactivated over processing time (King & Dehaene, 2014). Statistical significance of classification accuracies against chance (right-tail test against the chance level of 50%) and of the paired differences in accuracy between within- and across-domain classifications (two-tail test against an average accuracy difference of 0%) were established using sign permutation tests with cluster-based correction for multiple comparisons over time-points (cluster-defining threshold p-value = 0.05, alpha = 0.05; similar to the procedure of e.g. Cichy, Pantazis, & Oliva (2014).

### 2.8. Group representational dissimilarity analyses (RDMs)

We examined group RDMs (Cichy et al., 2014; Groen et al., 2012; Guggenmos, Sterzer, & Cichy, 2018; Kaneshiro et al., 2015; Kietzmann et al., 2017; Kong et al., 2020), comprised of average pairwise classification accuracy for each possible pair of visual images. For each group-average RDM, a split-half noise ceiling was also estimated, indicating the highest expected correlation given the level of noise in the data (e.g., Nili et al., 2014). Pearson’s correlations between group average RDMs were computed from random half-splits of the group data (i.e., half of the participants in each split), correcting these split-half correlations using the square root of the Spearman-Brown formula (Lage-Castellanos, Valente, Formisano, & De Martino, 2019) and averaging these estimates over 100 random half-splits of the group data. Following the procedure described in Lage-Castellanos et al. (2019), the noise ceiling was defined to be zero (i.e., at chance) for negative split-half correlations. The statistical significance of the resulting noise ceiling estimate against chance was evaluated using one-sided empirical *p*-values derived from null distributions obtained from 10,000 null split-half noise ceiling estimates. Each of these 10,000 null split-half noise ceiling estimates was computed using the procedure described above, with one of the two group-average RDMs scrambled in each of 100 random half-splits of the group data. Similarities between group RDMs were computed using Pearson’s correlations. Significance was evaluated based on two-sided parametric *p*-values associated with these correlation coefficients. All *p*-values corresponding to correlations between different group RDMs (**Figure 5**, lower triangle values) and to noise-ceiling estimates for each RDM (**Figure 5**, diagonal values) were corrected for multiple comparisons at the FDR level over this entire set of 21 correlations and noise-ceiling *p*-values.

### 2.9. Comparison of group RDMs with extant computational models of vision

Based on the available rankings of Brain-Score (Schrimpf et al., 2018), we selected CORnet-S (Kubilius et al., 2018) and the pool3 layer of VGG-16 (Simonyan & Zisserman, 2015) as potential computational models of adult high-level (inferotemporal) and mid-level (V4) vision, respectively. At the time of selection, CORnet-S was the highest overall ranking model and one of the highest ranking models for matching high-level visual regions (inferotemporal cortex); the pool3 layer of VGG-16 was listed as the highest ranking model for matching the mid-level visual region of V4. For VGG-16, we used the Matlab implementation pretrained on ImageNet. For CORnet-S, we used the pretrained implementation openly available at https://github.com/dicarlolab/CORnet. For each model, we obtained model activations in response to each of the 8 visual images used in the human experiment, then used pairwise Pearson’s correlations to compute Representational Similarity Matrices (RSMs) based on these activations. We also constructed a control RSM of low-level similarity using the Matlab implementation of SSIM (Wang, Bovik, Sheikh, & Simoncelli, 2004), an index of low-level similarity between two images that ranges from −1 to +1, taking a value of +1 when the two images are identical. Three model RDMs (CORnet-S, VGG-16, and control SSIM) were derived from these three RSMs by taking RDM = 1 – RSM (Cichy et al., 2014). Because a linear relationship between model and human representational distances could not be assumed, model RDMs were compared to experimental human RDMs using Spearman rank-correlations (**Figure 6**).

### 2.10. Data and Code Availability

Datasets and code generated during this study are available at [Figshare DOI to be activated upon publication acceptance].

## 3. Results

### 3.1. Event-Related Potentials (ERPs)

To form a basis of comparison with prior ERP work in infants, we first examined the observed average ERPs in each condition (animals, human body parts) and group (infants, adults) using commonly used regions and time-windows of interest (**Figure 1**). We used one-way mixed-effects ANOVAs to estimate the effect of domain (animal, body) on the average amplitude of the N290 (occipital ROI, 190-300ms), P400 (occipitotemporal ROI, 300-500 ms), and NC (central ROI, 300-500 ms) in infants. The effect of domain was not significant for any of these components (all *p*s > 0.15). A one-way mixed-effects ANOVA further examined the effect of categorical domain (animal, body) on the average amplitude of the adults’ N170 component (T5/P7 and T6/P8; 150–190 ms; e.g. Balas & Koldewyn, 2013). There was a significant effect of domain on the adults’ N170 (*F*[1,14] = 12.64, *p* = 0.003, 95%CI [-5.63; −1.39], d = −0.87), with more negative N170 amplitudes in response to human body than to animal images. Cluster-corrected comparisons of average waveforms at each ROI, unrestricted to time-windows of interest, yielded similar results (**Figure 1**), with statistically significant effects of domain (body vs. animal) in ROIs in adults, and no or statistically marginal effects in infants.

**Figure 1.**
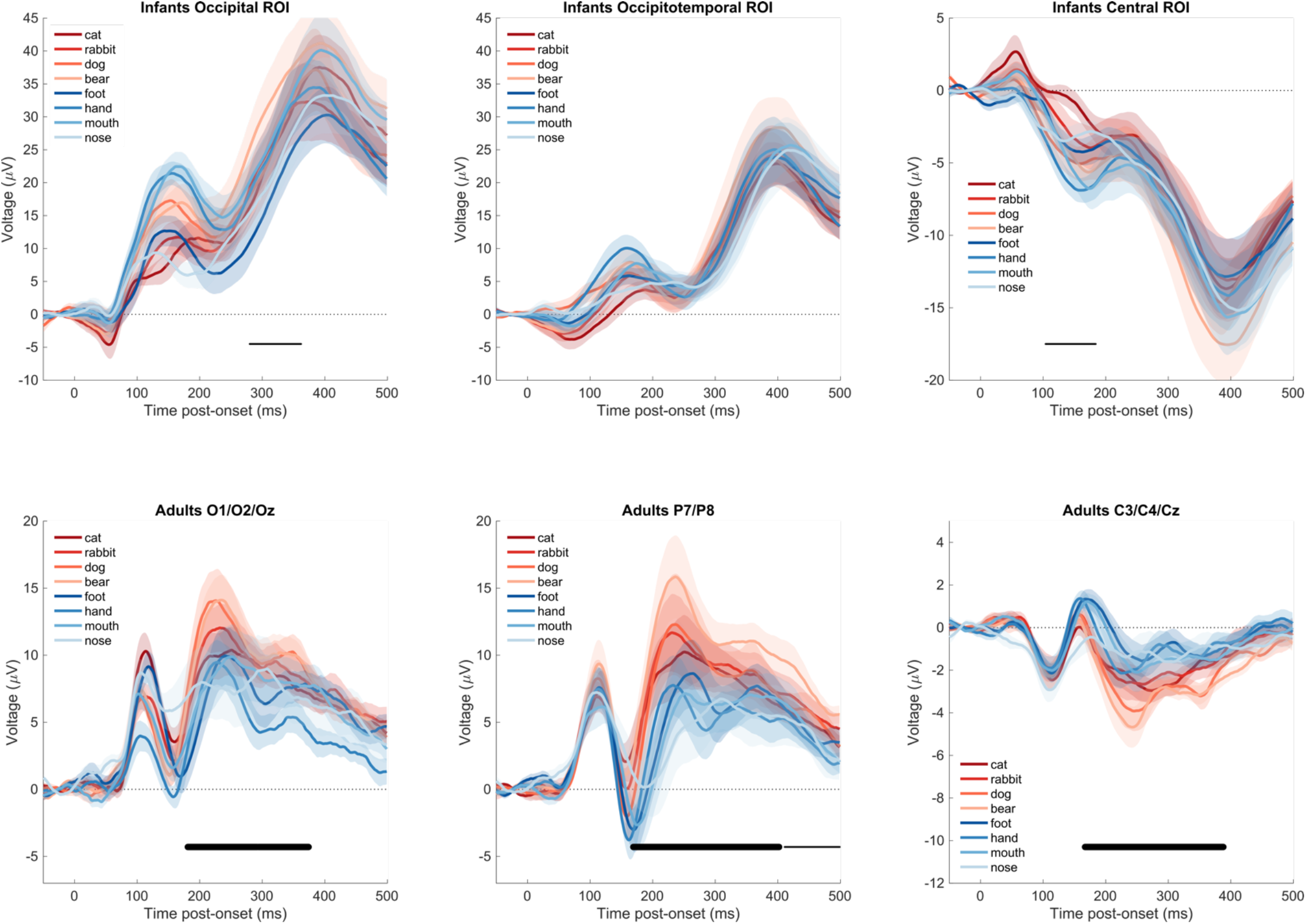
Average ERP waveforms in response to the visual stimuli in infants (12-15-month-olds) and young adults. Average ERP waveforms ± s.e.m, computed from the same data epochs that were used to perform within-subjects classification. In contrast to within-subjects classification, amplitudes were not z-scored, and were averaged over electrodes of interest. Thick (resp. thin) black horizontal lines indicate significant (resp. marginally significant) clusters for the difference in average amplitude in response to animals vs. body items (two-tailed).

### 3.2. Multivariate classification timeseries

We next conducted time-resolved multivariate classification of the EEG data (Cichy et al., 2014; Groen et al., 2012; Grootswagers et al., 2017; Isik et al., 2014; Kaneshiro et al., 2015; Kietzmann et al., 2017), representing the time-course of information available in the measured neural signals that identifies which of the 8 visual stimuli has been presented on a given trial. Specifically, a linear support vector machine (SVM) algorithm was used to classify trials as containing each member of all possible pairs of visual stimuli (e.g., cat versus dog) and at each millisecond time-point after stimulus onset. Neural representations could accurately discriminate between the presented visual stimuli in both infants and adults, averaging over all pairwise classifications (**Figure 2**, left column; right-tail comparison against a chance level of 50%, cluster-corrected sign permutation tests, cluster-defining threshold *p* < 0.05, corrected significance level *p* < 0.05). Results in adults replicated previous work (e.g. Cichy et al., 2014; Isik et al., 2014) showing sustained, above-chance average pairwise classification of visual images emerging by 100 ms and peaking at 72.86% classification accuracy at 177 ms (significant cluster from 82-499 ms). In infants, classification of visual images also rose to above-chance levels by 100 ms but peaked at 57.13% classification accuracy and about 150 ms later than in adults at 320 ms post-onset (significant clusters from 83-198 ms and from 212-361ms; additional marginally significant cluster from 419-499 ms). Using multi-class classification instead of average pairwise classification yielded comparable results (**Supplementary Figure S2**). Multivariate neural patterns differentiating between all 8 visual images (Haufe et al., 2014) are shown in **Figure 3**, exhibiting the expected occipital topography in infants, and occipitotemporal topography in adults.

**Figure 2.**
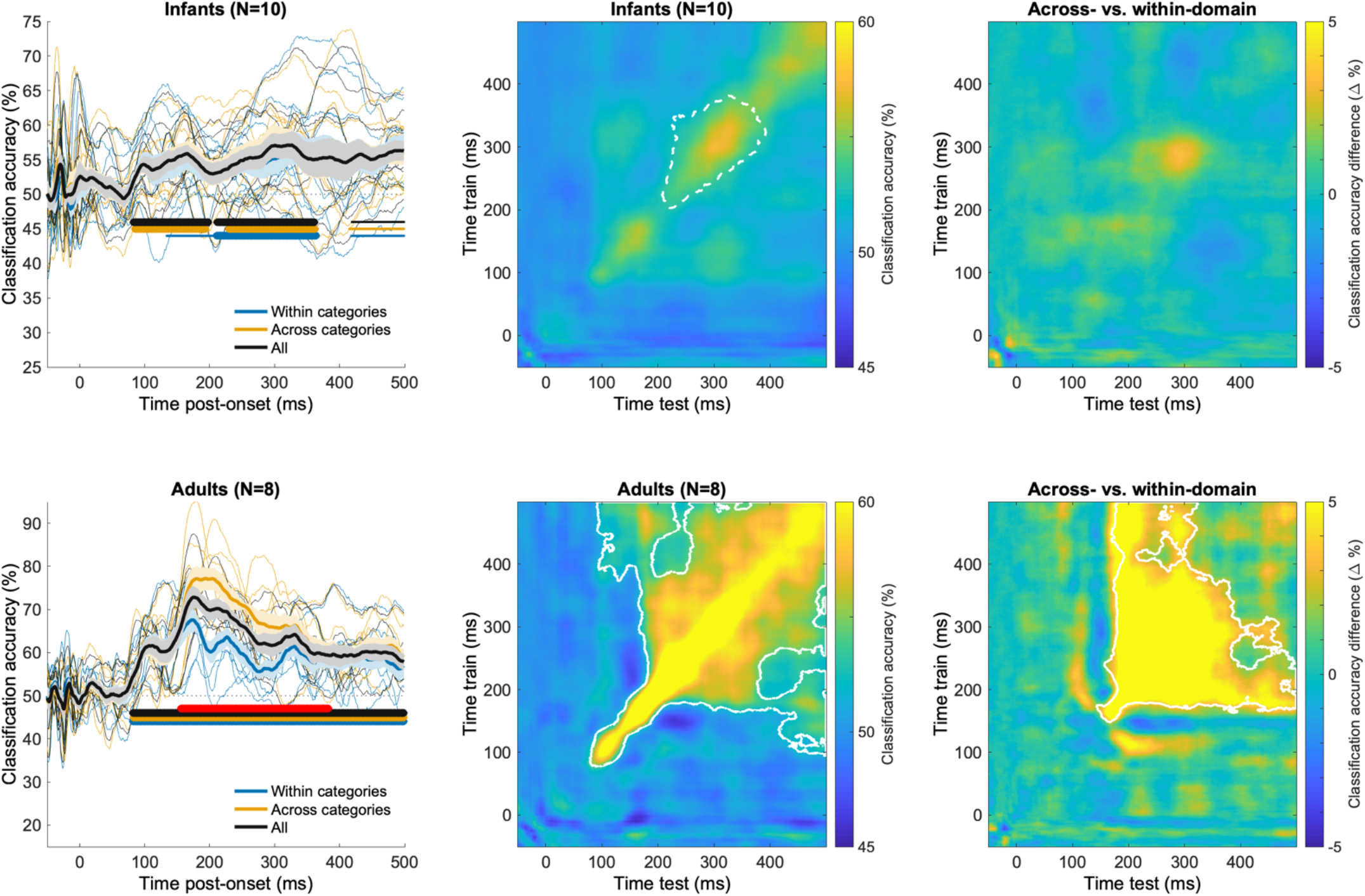
Average accuracy for the pairwise classification of visual stimuli in infants (12-15-month-olds) and adults. **Left:** Average ± s.e.m. accuracy timeseries for pairwise, within-subject classification within (e.g. cat vs. dog) or across domains (e.g. cat vs. hand). Thick (resp. thin) horizontal lines indicate statistically significant (resp. marginally significant) clusters of the difference between accuracy and the chance level of 50% (one-tailed). Red horizontal lines indicate significant clusters for the difference in accuracy between classifications across vs. within domains (one-tailed). **Middle:** Average, within-subject time generalization accuracy for all pairwise classifications. Solid (resp. dotted) white lines indicate the border of statistically (resp. marginally) significant 2-D clusters of the difference between accuracy and the chance level of 50%. **Right:** Average difference of within-subject time generalization accuracy for pairwise classifications across (e.g. cat vs. hand) vs. within (e.g. cat vs. dog) domains (e.g. cat vs. dog). White lines indicate the border of statistically significant clusters of this difference. See also **Supplementary Figure 2**

**Figure 3.**
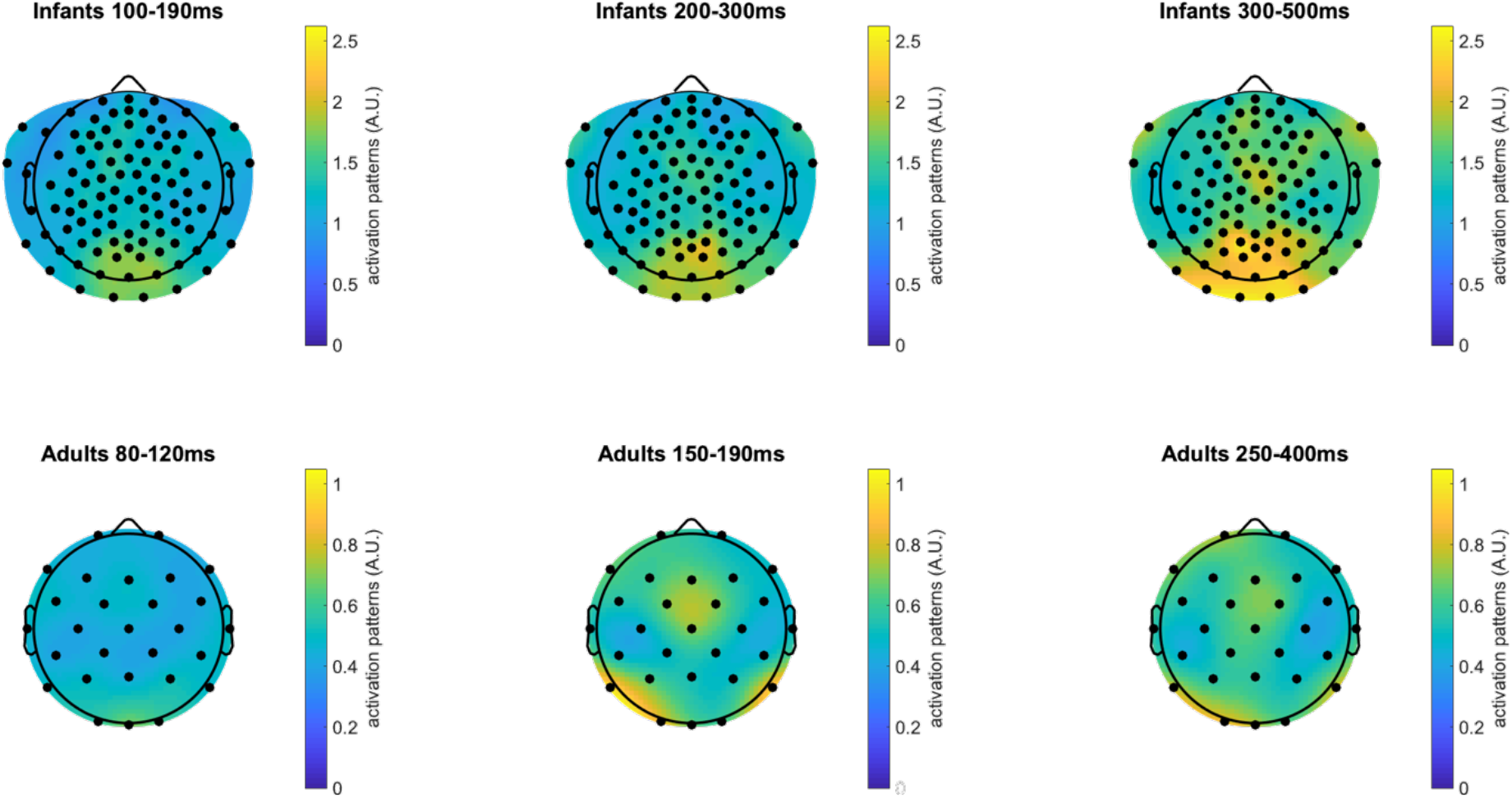
Multivariate channel patterns supporting pairwise classification of visual stimuli in infants (12-15-month-olds) and young adults. Average absolute activation patterns (Haufe et al, 2014) for pairwise, within-subject classification of visual stimuli (e.g. cat vs. dog, cat vs. hand, etc.). This method uses classifier weights and channel covariance to highlight channels were activation differences are contrastive between the stimulus classes.

To test whether neural representations discriminated pairs of visual images more strongly when those depicted objects came from a different categorical domain (e.g., dog versus hand) than when those depicted objects came from the same domain (e.g., dog versus cat), we next compared classification accuracies across and within domains (**Figure 2**, left column; two-tails comparison against the null hypothesis of 0% difference in accuracy, cluster-corrected sign permutation tests, cluster-defining threshold *p* < 0.05, corrected significance level *p* < 0.05). Results in adults replicated previous findings (Cichy et al., 2014), showing higher classification accuracy across than within domains by 200 ms post-onset (peak accuracy difference of +16.95% at 201 ms; significant cluster for the difference in accuracy from 156-383 ms). In infants, average pairwise accuracies for the across-domain and within-domain classifications did not differ significantly (cluster *p*s> 0.1).

Because infants had lower trial counts, their datasets could have failed to reveal a domain effect because of the overall lower number of trials per stimulus item. Thus, we asked if the domain effect that was evident in the adult dataset would remain if trial counts in the adult dataset were yoked to those of the infant dataset. When trial counts in the adult dataset were matched to those of the infant dataset, a domain effect remained significant (cluster *p* < 0.05) in the same direction and at the same overall timing as in the full adult dataset (**Supplementary Figure S3**). Overall, it appears unlikely that lower trial counts alone could account for the lack of a domain effect in the neural representations of infants. However, other differences between the adult and infant datasets (e.g., channel counts, attention, electrode type, cortical folding, skull thickness, etc.) could have played a role.

### 3.3. Temporal generalization

Robust neural representations can remain active for several tens of milliseconds, for example if the task requires the maintenance of that representation in working memory (Quentin et al., 2019). Temporal generalization analyses (King & Dehaene, 2014) were run to assess the maintenance of neural representations of visual images over time (**Figure 2**, central column; right-tail comparison against the chance level of 50%, cluster-corrected sign permutation tests, 2-D cluster-defining threshold *p* < 0.05, corrected significance level *p* < 0.05). Temporal generalization analyses quantify how well a classifier trained on neural data at a given time point (e.g., 100 ms post-onset) can decode neural data from the same participant at another time points (e.g. 150 ms post-onset), repeatedly for all time points available for training and testing, providing a metric of the consistency of neural representations across distinct time points (King & Dehaene, 2014). In adults, these analyses showed the maintenance (generalization) of neural representations supporting pairwise classification of stimuli along the temporal diagonal (i.e. for test time-points closely bordering the times at which the classifier was trained), as previously observed (e.g. Isik et al., 2014; King & Dehaene, 2014). Similar results were found in infants, but classification accuracy for generalizing across time-points was only marginally significant (2-D cluster *p* < 0.10). Thus, neural representations that support the classification of visual images may be maintained over time to some degree by 12-15-months of age, but not robustly enough to reach statistical significance. Because there were no task demands (i.e., passive viewing), it is possible that adults spontaneously held neural representations in working memory whereas infants did not, although perhaps they could if the task demanded it (e.g., delayed match-to-sample).

To test whether neural representations that support the differentiation of categorical domains were maintained over time, we next examined the difference in temporal generalization accuracy for classifying across-domain versus within-domain pairs of stimuli (**Figure 2**, right column; two-tails comparison against the null hypothesis of 0% difference, cluster-corrected sign permutation tests, 2-D cluster-defining threshold *p* < 0.05, corrected significance level *p* < 0.05). In adults, the average pairwise temporal generalization accuracy was significantly higher for across-than for within-domain classifications from roughly 150 ms post-onset (2-D cluster *p* < 0.05). Thus, in adults, the previously described domain organization of neural representations is dynamically maintained over processing time. In infants, we again found no significant difference in average decoding accuracy for across-versus within-domain classifications, mirroring the timeseries results (2-D cluster *p*s > 0.10).

### 3.4. Representational Dissimilarity Matrices

The foregoing results confirmed that neural representations in infants and adults supported the reliable decoding of visual stimuli, averaging over pairs of visual stimuli, and that there was some indication that these neural representations were maintained overtime. We next asked whether the underlying neural representations of each of these visual stimuli were consistent (a) across individuals of the same age group, (b) across time post-stimulus (i.e., between different temporal windows), (c) across age groups, and (d) with the way in which stimulus similarities are defined by computer vision algorithms. A standard way of visualizing the dynamic geometry of visual representations is to compute a Representational Dissimilarity Matrix (RDM), which consists of the average pairwise classification accuracy for each possible pair of visual stimuli in each age group, over four broad time-windows (**Figure 4**) defined a priori from visual ERP studies in infants and adults. As expected from existing work in adults (e.g. Cichy et al., 2014), RDMs in adults exhibited a clear organization by domain, with higher average pairwise accuracy across than within domains in the 150-190 ms and 250-400 ms, but not 80-120 ms time-windows (two-tail paired *t*-test, FDR-corrected; 80-120 ms: t[7] = − 0.54, *p* = 0.608; 150-190 ms: t[7] = 4.03, *p* = 0.008; 250-400 ms: t[7] = 7.50, *p* = 0.004). No such pattern was evident in the infants’ RDMs (all *p*s > 0.5).

**Figure 4.**
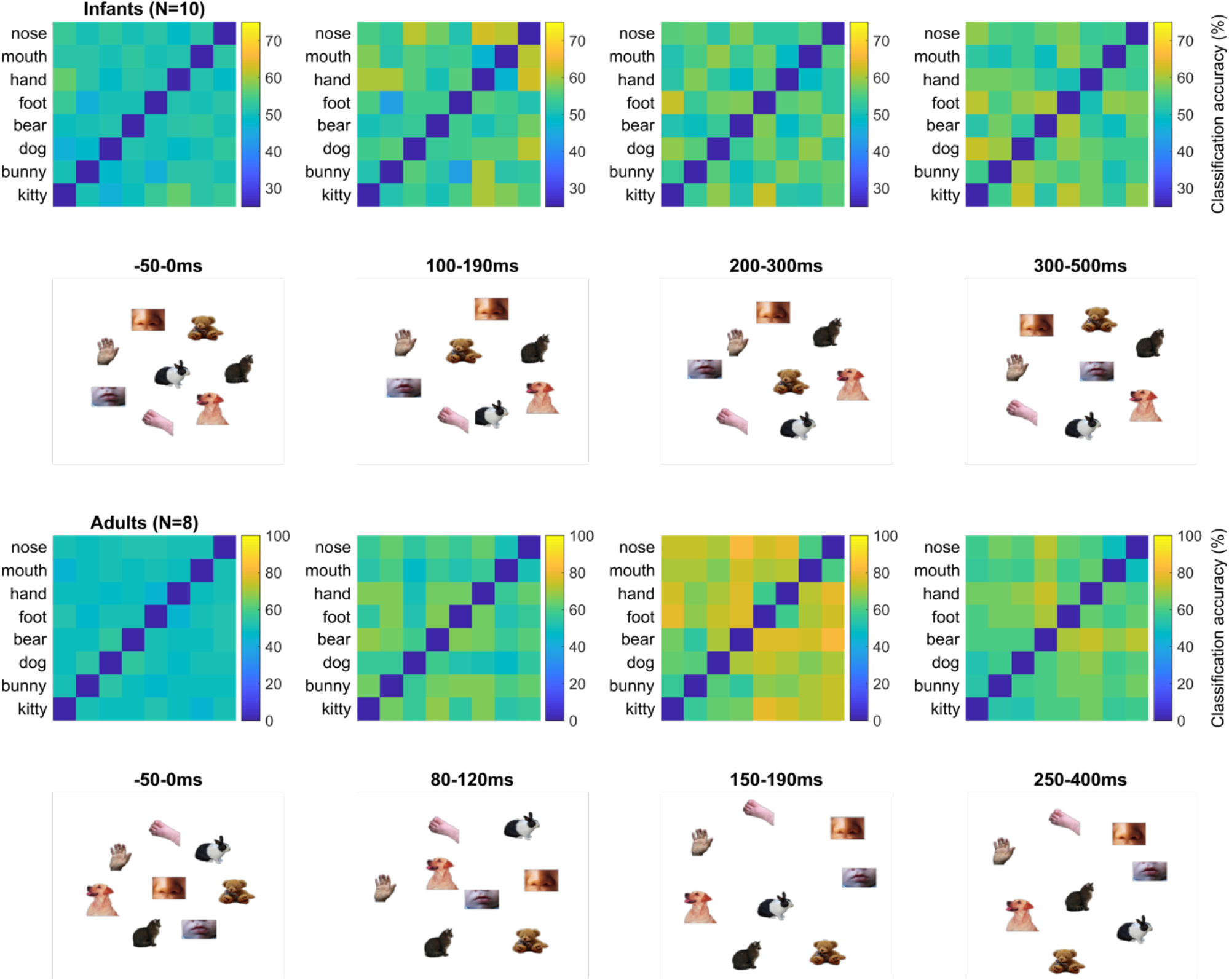
Representational dissimilarity of visual stimuli in infants (12-15-month-olds) and young adults. Group average accuracy for all pairwise, within-subject classifications, and 2D multidimensional scaling visualization (metric stress criterion).

#### 3.4.1 Group representational similarity analyses

Although the organization of infants’ neural representations may not exhibit a categorical boundary between animals and human body parts like adults do, infants may nevertheless have a reliable neural representation, but one that is organized differently from that of adults. That is, neural representations for distinguishing between animal and body images may not exhibit a linear domain boundary between animal and body items in infants but nevertheless exhibit a reliable organization that is similar amongst individuals of the same age. To investigate this possibility, we estimated the reliability of representational spaces measured at the group level. Specifically, we estimated the split-half noise-ceiling of each group-average using the upper half of the RDM (**Figure 5**, left panel, diagonal values) and compared it to an empirical chance level (see **Materials & Methods**). Pearson’s correlations between different group-average upper RDMs were additionally computed to assess representational similarity between different age groups, and between different time-windows within each age group (**Figure 5**, left panel, lower triangle and inset).

**Figure 5.**
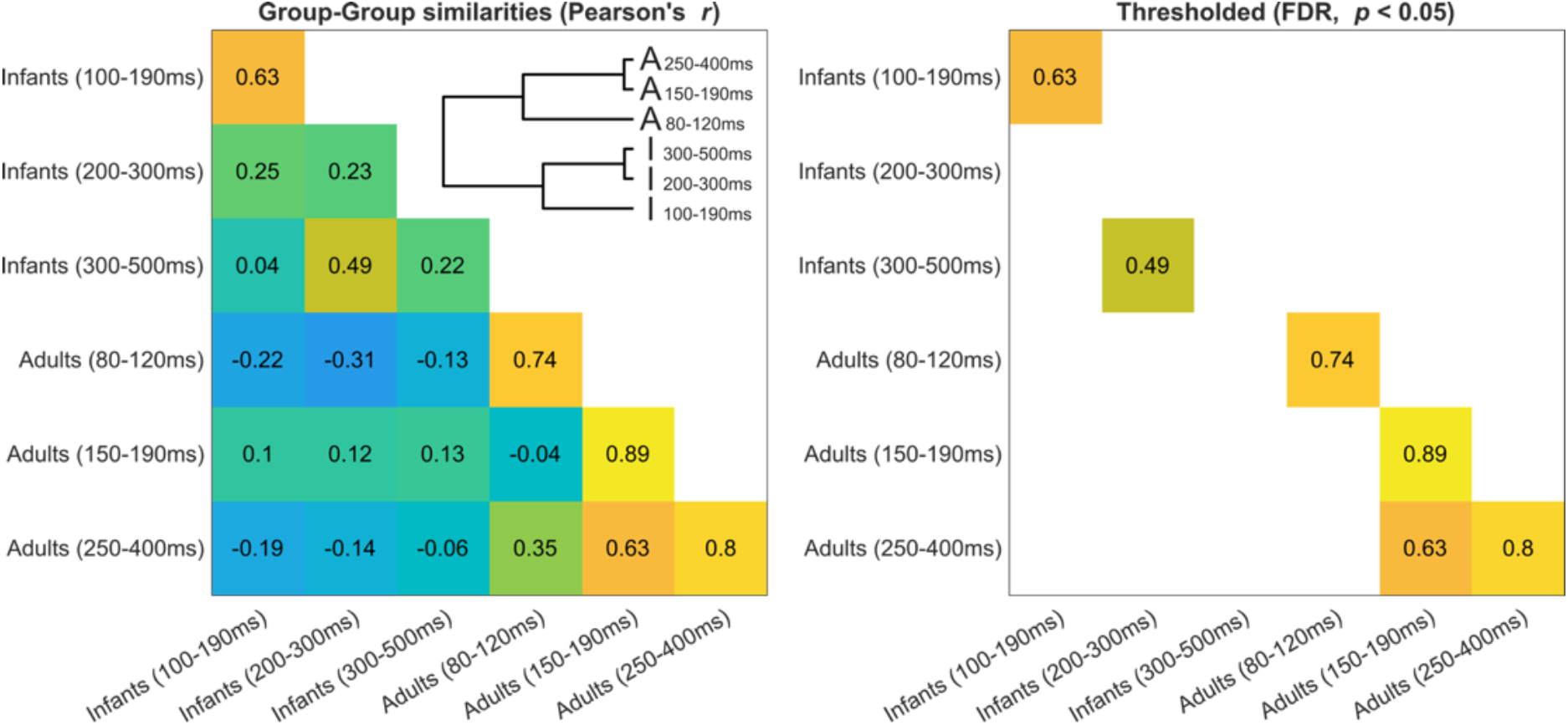
Group representational similarity. Lower triangle: Pearson’s correlations between group-average Representational Dissimilarity Matrices, as a function of time-window and age group. **Inset:** Dendrogram representation of correlations between group RDMs. **Diagonal:** Split-half noise ceiling estimates.

After FDR-correction for multiple comparisons, (**Figure 5**, right panel, diagonal values), RDM noise ceilings exceeded the empirical chance level in each time window in adults (80-120 ms: ρ_SHnc_ = 0.74, FDR-corrected *p* < 0.001; 150-190 ms: ρ_SHnc_ = 0.89, FDR-corrected *p* < 0.001; 250-400 ms: ρ_SHnc_ = 0.80, FDR-corrected *p* < 0.001) and one of the three time-windows in infants (100-190 ms: ρ_SHnc_ = 0.63, FDR-corrected *p* < 0.001). Thus, average dissimilarities of neural representations between pairs of visual images could reliably be estimated at the group level in these time-windows and were similar amongst distinct groups of individuals of the same age. Noise ceilings did not exceed the empirical chance level in the remaining two time-windows in infants (FDR-corrected *p*s > 0.9), suggesting that these group representational dissimilarity spaces could not be reliably estimated in this time-windows – likely due to limitations in sample size or increased heterogeneity among participants.

We next asked whether group RDMs were similar across different age groups, or across different time-windows within each age-group. After FDR-correction for multiple comparisons, no significant positive correlation between RDMs from different age groups were found (FDR-corrected *p*s > 0.2; **Figure 5**, right panel, lower triangle values). In adults, the group average RDMs from the 150-190 ms and 250-400 ms time-windows were significantly correlated (ρ = 0.63, FDR-corrected *p* = 0.001; **Figure 5**, right panel, lower triangle values), as could be expected from the documented maintenance of neural representations for discriminating visual images over this time-frame in adults (**Figure 2**, bottom row). Similarly, in infants, the group average RDMs from the 200-300 ms and 300-500 ms time-windows were also significantly correlated (ρ = 0.49, FDR-corrected *p* = 0.026; **Figure 5**, right panel, lower triangle values). Overall, within each age group, group RDMs corresponding to the two later time-windows (200-300 and 300-500 ms in infants 150-190 and 250-400 ms in adults) were positively correlated with one another but not with the group RDM corresponding to the earlier time-window (100-190 ms in infants or 80-120 ms in adults); group RDMs did not correlate above chance levels between age groups.

#### 3.4.2. Comparison with models of vision

Because at least some of the group-level RDMs could be reliably estimated, we next used Spearman’s rank-correlations to ask whether any group-level RDMs shared similarities with the representational geometries predicted by two of the current best models of object vision (Schrimpf et al., 2018), CORnet-S (Kubilius et al., 2018) and VGG-16 (Simonyan & Zisserman, 2015), or by a control measure of low-level image similarity (SSIM, Wang, Bovik, Sheikh, & Simoncelli, 2004). Two exploratory correlations passed the uncorrected threshold of statistical significance: a positive correlation between the CORnet-S RDM and the adult 80-120 ms RDM (Spearman *r* = 0.49, *p* = 0.010), and a negative correlation between the adult 250-400 ms RDM and the control SSIM RDM (Spearman *r* = −0.47, *p* = 0.012; **Figure 6**). None survived FDR-correction for multiple comparisons over the entire set of 18 Spearman’s rank-correlations tested, and no other significant correlations were found (**Figure 6**).

**Figure 6.**
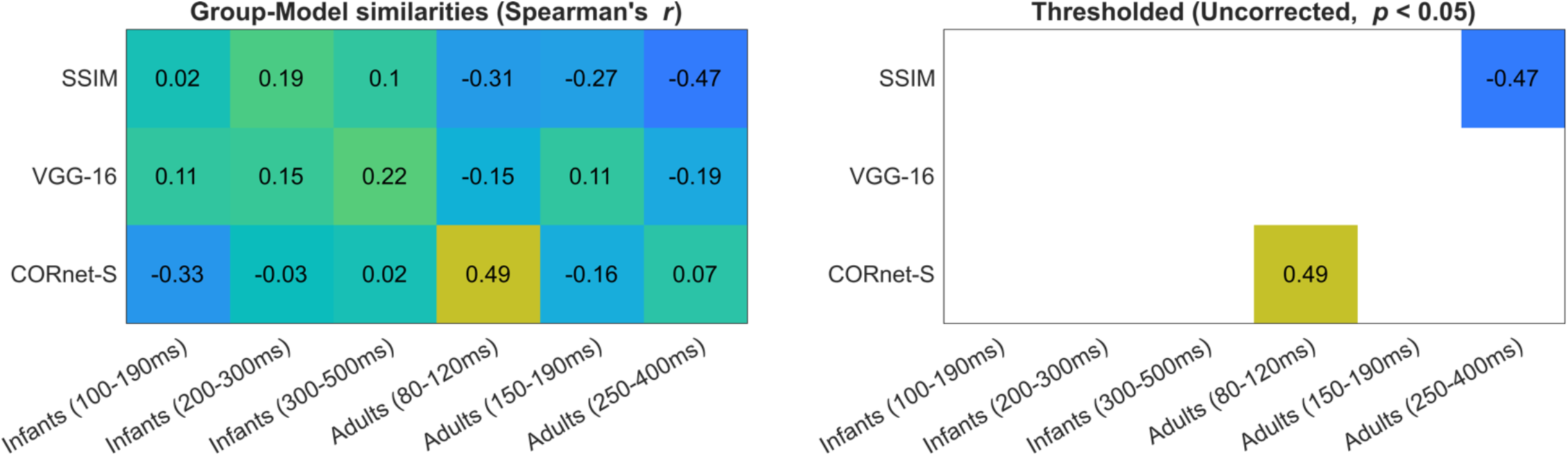
Group representational similarity with models of vision. Similarities between group Representational Dissimilarity Matrices and Representational Dissimilarity Matrices derived from three models: the final layer of CORnet-S (computational model of high-level vision), the pool3 layer of VGG-16 (computational model of mid-level vision), and a low-level control index (SSIM). Similarities that passed an uncorrected significance threshold (uncorrected p < 0.05) are indicated on the right-side panel; none remained significant after FDR-correction.

## 4. Discussion

In the present study we provide a proof of concept for employing time-resolved multivariate pattern analysis methods with EEG data to characterize the dynamics of neural representations of visual stimuli in infants, and to characterize group-level representational spaces across infants and adults. Neural representations of visual stimuli in infants support the reliable classification of 8 different visual images but appear to be only modestly sustained over the post-stimulus time period and do not exhibit the robust differentiation between the domains of animals versus human body parts that is evident in adults. Group-average representational spaces, as indexed by Representational Dissimilarity Matrices based on pairwise classification accuracy, were found to meet a basic standard of reliability in some but not all cases. Overall, group average representational spaces appear similar across some time-windows within each age group, but not across age groups.

To our knowledge, the current study is the first to investigate the dynamics of neural representations of preverbal infants using a decoding framework. Neural representations supported the classification of visual images in 12-15-month-olds. The feasibility of estimating encoding models that directly quantify the association between multiple stimulus features and changes in infant EEG data has been demonstrated in the case of low-level sensory features (such as luminance or amount of motion) in audiovisual cartoon movies, suggesting that the approach may allow for building encoding models corresponding to more complex stimulus dimensions (Jessen, Fiedler, Münte, & Obleser, 2019). We extend these results by demonstrating the feasibility of pairwise decoding of visual stimuli in 12-15-month-olds. Together with infant fNIRS decoding (Emberson et al., 2017), infant fMRI representation similarity analysis (Deen et al., 2017), and infant EEG encoding models (Jessen et al., 2019), the current findings contribute to the utilization of information-based, multivariate, computational methods as a powerful toolkit for analyzing infant neural data.

Evidence for the maintenance of neural representations of visual images over time within the first 500 ms of processing was weak in 12-15-month-olds. It is possible that stronger evidence would have been found for the maintenance of representations beyond the first 500 ms of processing. For example, non-linear increases in response to supraliminal visual stimuli, potential neural markers of conscious access to these stimuli, emerge from roughly 750 ms post-onset in 12-15-month-olds versus from roughly 300 ms in adults (Kouider et al., 2013). Because the maintenance and reactivation of sensory representations over-time is thought to be characteristic of conscious access, it is conceivable that the maintenance of visual representations over-time could be evident later than 500 ms in 12-15-month-olds. Future research may address this question by utilizing paradigms that allow for the analysis of later responses (after 500 ms) in this age range while preserving a high enough number of artefact-free trials for decoding.

We examined whether decoding accuracies would be higher, overall, when classifying trials according to pairs of stimuli that belong to a different domain (animal versus parts of the human body) compared to those that came from the same domain, as had previously been reported in adults (e.g., Cichy et al., 2014). This pattern of results was evident in adults but absent in infants. This negative finding suggests that the visual cortex of infants may not represent visual images of animal and human bodies in a manner that linearly, robustly differentiates their categorical domain (i.e., through differences in neural activations that are consistent in pattern and timing across multiple trials). Reducing trial numbers in the adult dataset to those of the infant dataset did not eliminate the domain effect in adult classification accuracies, suggesting that factors beyond trial numbers were responsible for these null findings in infants. Converging results were found when estimating the effect of domain on univariate ERP components, with a clear effect present in adults but less so in infants. Taken together, the current EEG findings are aligned with the previous fMRI findings of Deen et al., (2017), according to which some functional domain specificity of the visual cortex is already present by the end of the first year of life but is less (or differently) marked than in adults. Alternatively, neural representations that support differentiating animals from parts of the human body may be apparent in patterns of neural activity that were not considered in the current analysis – such as activity beyond 500 ms post-onset, or non-linear aspects of neural responses including evoked and induced oscillations. An effect of category on multivariate classification accuracy may have been obtained in infants for domain categories of visual images for which univariate differences have been consistently found, such as faces vs. objects or human vs. non-human faces (de Haan et al., 2002; Farzin et al., 2012; Guy, Zieber, & Richards, 2016; Peykarjou et al., 2014, 2017), or in a paradigm designed to induce or reveal categorization via a novelty detection response (e.g., oddball; Marinović et al., 2014). Factors beyond trial counts (such as signal quality, EEG equipment brand, stimulus duration, skull thickness, cortical folding, etc.) may also have contributed to the lack of a domain effect in the infant data. The absence of a category domain effect in the current infant data likely reflects both the current methodological approach (which was limited to examining time-locked, linear patterns of EEG voltage amplitude across trials in a small sample of infants) and meaningful functional differences in the way that infants represent visual objects compared to adults. That is, it is possible that a different paradigm or measure may have elicited a linear category boundary for human body vs. animal images in 12-15-month-olds. Even then, however, the absence of such a linear boundary in the current data, if reproducible in larger samples and not attributable to trivial measurement differences (e.g., skull thickness), would be consistent with the notion that visual cortex undergoes a developmental change in the *manner by which* animal vs. human body images become “untangled” (DiCarlo & Cox, 2007) during visual processing between infancy and adulthood. Future research may uncover the mechanisms by which the neural representations of visual objects become organized along categorical domains by adulthood and clarify the association between the increasingly specific parcellation of functional domains within the visual cortex and the representation of domains these networks support.

We explored the group-level Representational Dissimilarity Matrices (RDMs) implied by average pairwise classification accuracies for each possible pair of visual stimuli presented. No similarity was found between RDMs across age groups, although some similarities were found between RDMs in different time-windows within age groups. The results aligned with those of Deen et al., (2017), who reported that fMRI-derived RDMs were similar within the infant group and within the adult group, but dissimilar between these two age groups. The noise ceiling of the current results was generally low for infant RDMs, as several group-level RDMs could not be reliably estimated based on the current datasets. Thus, the current findings likely underestimate the extent to which group-level infant representational spaces may linearly discriminate visual domains, resemble other group-level representational spaces within or across age groups, or resemble representational spaces derived from model algorithms.

The current study provides a first proof-of-concept for the use of decoding analyses from infant EEG signals. The high attrition rate and subsequently small sample sizes in the infant group limits the extent to which the current results may generalize, and likely limited the reliability with which group-level RDMs could be estimated. Future research utilizing similar methods should attempt to decrease attrition by increasing the total number of valid trials collected from each infant (e.g., through changes in paradigm or signal processing), or by adapting analysis methods to accommodate small numbers of trials per individual participant and stimulus condition. The relatively low number of individual stimuli (8) used limited the statistical power of analyses comparing different group-level RDMs or comparing these RDMs with models of vision. The stimulus set used did not attempt to overly correct for low-level visual differences between stimuli, nor did it attempt to elicit object-specific responses invariantly to changes in size, lightning, viewpoint, etc. Thus, pairwise classification accuracies were tracking the dynamics of neural representations that support differentiating between different visual *images* (a specific picture of a cat versus a specific picture of a dog), as opposed to differentiating between different *objects* per se (multiple exemplars of cats and dogs). Future research may examine whether neural representations of visual objects generalize across changes in size, view, or other dimensions infants as they do in adults (e.g. Isik et al., 2014).

## 5. Conclusions

In conclusion, we demonstrate the feasibility of time-resolved multivariate pattern analysis methods with infant EEG data at the beginning of the second postnatal year, and a first characterization of the dynamics of multivariate neural representations supporting the differentiation of animate visual stimuli in the infant brain. Univariate (activation-based) and multivariate (information-based) analyses converged to suggest that the categorical domain differentiation of neural responses to and representations of visual stimuli depicting animals versus parts of the body during passive viewing, robustly observed in adults and thought to reflect functional organization in the visual cortex, is not fully in place by 12-to 15-months of age. Future research will determine if this conclusion extends to other domains of visual stimuli and illuminate developmental mechanisms that lead to such differentiations by adulthood.

## Acknowledgements

This work was funded by NSF EAGER-1514351 to RNA and CAN, a grant and an Emmy Noether Grant (CI-241/1-1) to RMC, and a Philippe Foundation award to LB. We thank the participants, families, and students who made this work possible, Dr. Dimitrios Pantazis for sharing MATLAB code to perform cluster-based inferences on classification time-series, and Heather Kosakowski and Kirsten Lydic for helpful feedback on an earlier version of this manuscript.

## Author Contributions

Conceptualization, Funding Acquisition, & Supervision, R.N.A., C.A.N, and R.M.C.; Methodology, R.N.A., C.A.N, R.M.C., B.Z., and L.B; Software L.B., R.M.C., B.Z.; Formal Analysis & Visualization, L.B.; Investigation & Data Curation, E.R., J.C., Z.P., and L.B.; Writing – Original Draft, L.B., C.A.N., and R.N.A‥; Writing ‒Review & Editing, B.Z., R.M.C.;, R.N.A, C.A.N., and R.M.C.

