## Supplementary Materials for "Temporal dynamics of visual representations in the infant brain"

### Supplementary Figures and Tables

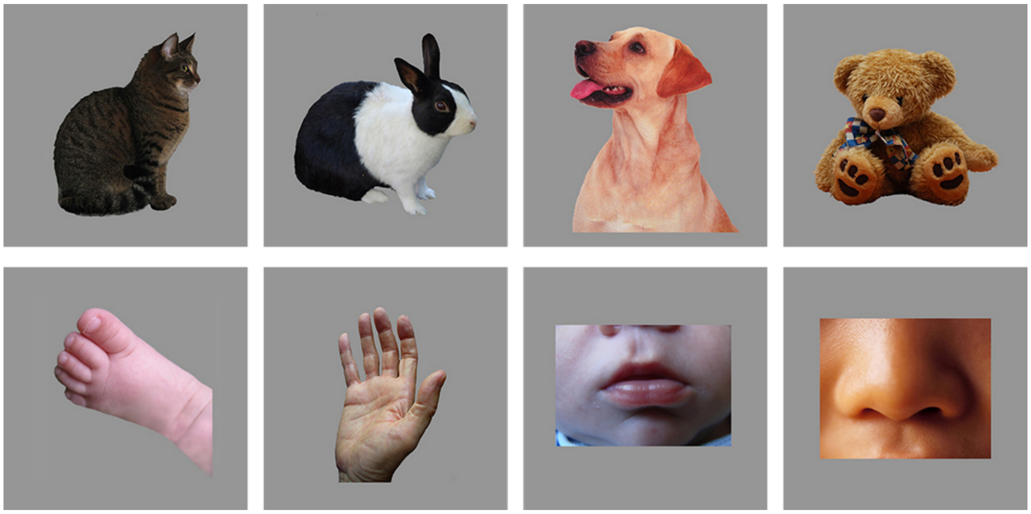

**Supplementary Figure S1. Visual stimuli.** Animate visual stimuli consisted in 8 pictures of animals (cat, rabbit, dog, and teddy bear) and parts of the human body (hand, foot, nose, mouth).

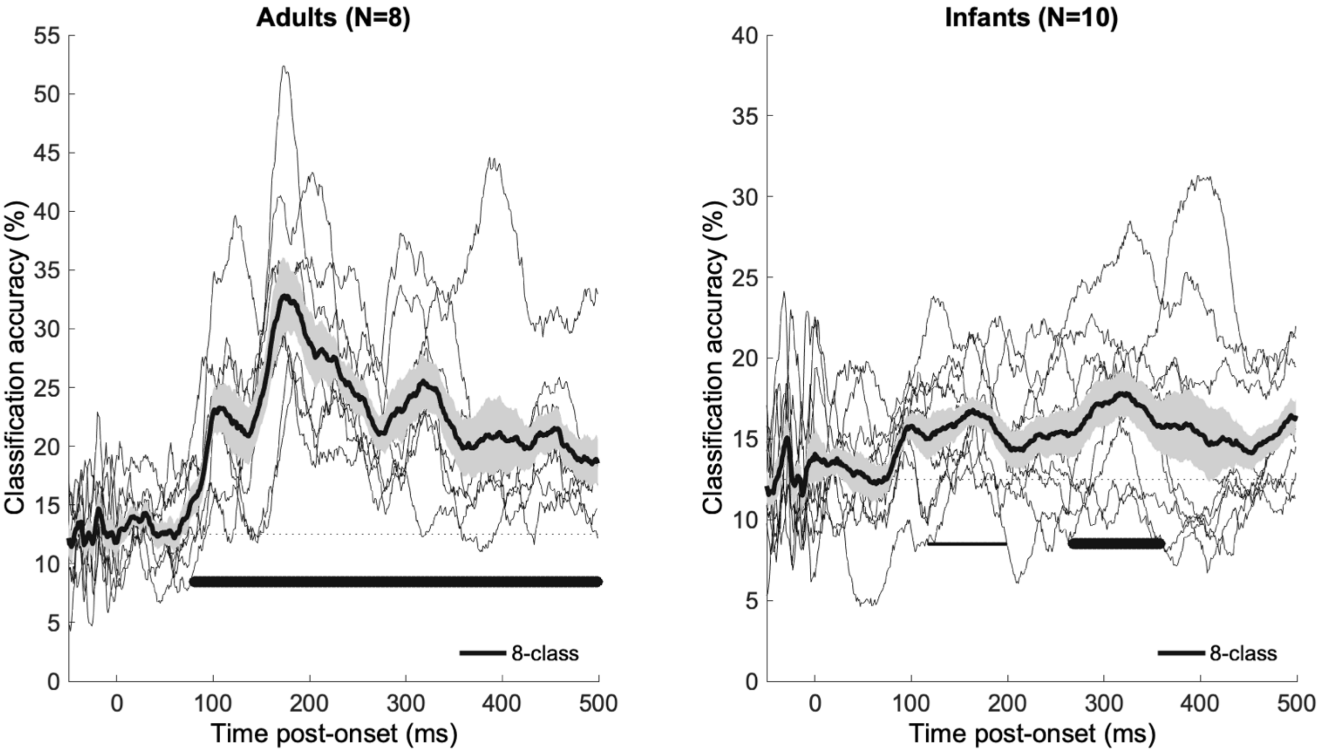


**Supplementary Figure S2. Average accuracy for the multiclass classification of visual stimuli in infants (12-15-month-olds) and adults.** Average ± s.e.m. accuracy timeseries for 8-class, within-subject classification. Thick (resp. thin) horizontal lines indicate statistically significant (resp. marginally significant) clusters of the difference between accuracy and the chance level of 12.5% (one-tailed).

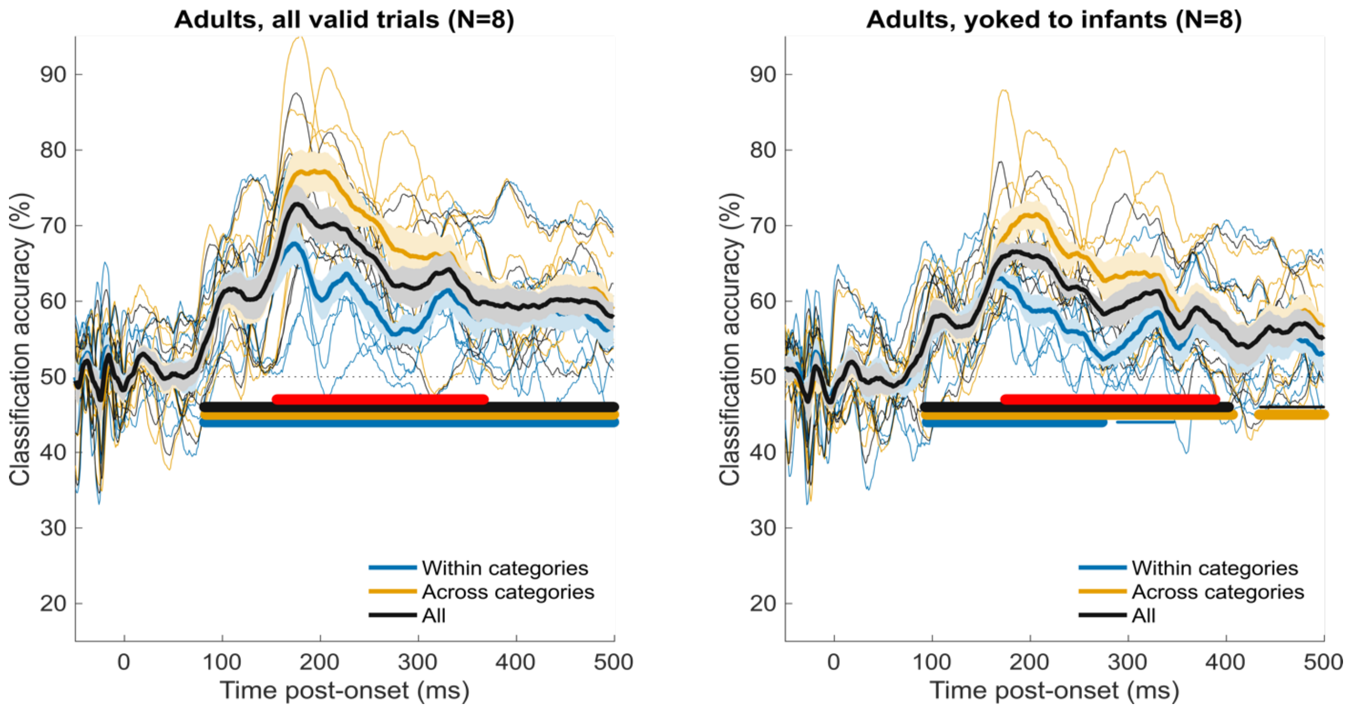


**Supplementary Figure S3. Average accuracy for the pairwise classification of visual stimuli in young adults, estimated from all valid trials (left) or valid trial counts yoked to the infant dataset (right).** Average ± s.e.m. accuracy timeseries for pairwise, within-subject classification within (e.g. cat vs. dog) or across categories (e.g. cat vs. hand). Thick (resp. thin) horizontal lines indicate statistically significant (resp. marginally significant) clusters of the difference between accuracy and the chance level of 50% (one-tailed). Thick (resp. thin) red horizontal lines indicate significant (resp. marginally significant) clusters for the difference in accuracy between classifications across vs. within categories (one-tailed). In the adults' dataset yoked to infants, participant and/or trials were discarded in reverse chronological until numbers of participants and valid trials per participant and condition were exactly equal to (or lower than) that of the infant dataset.

Supplementary Table 1. Summary data from the MacArthur Communicative Development Inventories: Words and Gestures (CDI).

| | | | 12-15-month-old infants |
|---|---|---|---|
| Total N of subsample with CDI data | | | 15 |
| Also included in EEG analyses | | | 7 |
| CDI raw scores (Mean ± SD) | | | |
| Words | | | |
| | Words understood (Max: 396) | | 107.47 ± 64.94 |
| | Words produced (Max: 396) | | 10.60 ± 8.02 |
| | Total (Max: 792) | | 118.07 ± 67.01 |
| Gestures | | | |
| | Early gestures (Max: 18) | | 12.87 ± 2.29 |
| | Late gestures (Max: 45) | | 19.60 ± 7.91 |
| | Total (Max: 63) | | 32.47 ± 9.16 |
| CDI individual word items (% Infants) | | | |
| Understands | | | |
| | Animals | bear | 20.00% |
| | | bunny | 26.67% |
| | | cat | 46.67% |
| | | dog | 93.33% |
| | | kitty | 26.67% |
| | | teddy bear | 20.00% |
| | Body parts | foot | 46.67% |
| | | hand | 40.00% |
| | | mouth | 53.33% |
| | | nose | 60.00% |
| | At least one of the above 10 words | | 100.00% |
| Produces | | | |
| | Animals | bear | 0.00% |
| | | bunny | 0.00% |
| | | cat | 6.67% |
| | | dog | 53.33% |
| | | kitty | 0.00% |
| | | teddy bear | 0.00% |
| | Body parts | foot | 0.00% |
| | | hand | 0.00% |
| | | mouth | 6.67% |
| | | nose | 0.00% |
| | At least one of the above 10 words | | 60.00% |
